# Evolving a mitigation of the stress response pathway to change the basic chemistry of life

**DOI:** 10.1101/2021.09.23.461486

**Authors:** Isabella Tolle, Stefan Oehm, Michael Georg Hoesl, Christin Treiber-Kleinke, Lauri Peil, Abdul-Rahman Adamu Bukari, Torsten Semmler, Juri Rappsilber, Aleeza Gerstein, Nediljko Budisa

## Abstract

Billions of years of evolution have produced only slight variations in the standard genetic code, and the number and identity of proteinogenic amino acids have remained mostly consistent throughout all three domains of life. These observations suggest a certain rigidity of the genetic code and prompt musings as to the origin and evolution of the code. Here we conducted an adaptive laboratory evolution (ALE) to push the limits of the code restriction, by evolving *Escherichia coli* to fully replace tryptophan, thought to be the latest addition to the genetic code, with the analog L-β-(thieno[3,2-*b*]pyrrolyl)alanine ([3,2]Tpa). We identified an overshooting of the stress response system to be the main inhibiting factor for limiting ancestral growth upon exposure to β-(thieno[3,2-*b*]pyrrole ([3,2]Tp), a metabolic precursor of [3,2]Tpa, and Trp limitation. During the ALE, *E. coli* was able to “calm down” its stress response machinery, thereby restoring growth. In particular, the inactivation of RpoS itself, the master regulon of the general stress response, was a key event during the adaptation. Knocking out the *rpoS* gene in the ancestral background independent of other changes conferred growth on [3,2]Tp. Our results add additional evidence that frozen regulatory constraints rather than a rigid protein translation apparatus are Life’s gatekeepers of the canonical amino acid repertoire. This information will not only enable us to design enhanced synthetic amino acid incorporation systems but may also shed light on a general biological mechanism trapping organismal configurations in a status quo.

**SIGNIFICANCE STATEMENT:** The (apparent) rigidity of the genetic code, as well as its universality, have long since ushered explorations into expanding the code with synthetic, new-to-nature building blocks and testing its boundaries. While nowadays even proteome-wide incorporation of synthetic amino acids has been reported on several occasions^1–3^, little is known about the underlying mechanisms.

We here report ALE with auxotrophic *E. coli* that yielded successful proteome-wide replacement of Trp by its synthetic analog [3,2]Tpa accompanied with the selection for loss of RpoS^4^ function. Such laboratory domestication of bacteria by the acquisition of *rpoS* mitigation mutations is beneficial not only to overcome the stress of nutrient (Trp) starvation but also to evolve the paths to use environmental xenobiotics (e.g. [3,2]Tp) as essential nutrients for growth.

We pose that regulatory constraints rather than a rigid and conserved protein translation apparatus are Life’s gatekeepers of the canonical amino acid repertoire (at least where close structural analogs are concerned). Our findings contribute a step towards understanding possible environmental causes of genetic changes and their relationship to evolution.

Our evolved strain affords a platform for homogenous protein labeling with [3,2]Tpa as well as for the production of biomolecules^5^, which are challenging to synthesize chemically. Top-down synthetic biology will also benefit greatly from breaking through the boundaries of the frozen bacterial genetic code, as this will enable us to begin creating synthetic cells capable to utilize an expanded range of substrates essential for life.

## INTRODUCTION

Evolution is an ongoing and pervasive process for all life on the planet. The mechanisms of evolution have yielded extant living organisms with immense biodiversity. Vast diversity exists along nearly all possible axes imaginable in body form, structure, and function. Given the potential for adaptation to explore different trajectories and organisms to live in virtually all known ecosystems on the planet, it is perhaps most unexpected not to find variation in the building blocks themselves: all life is based on the same 20 proteinogenic amino acids. The extant genetic code, which appeared >3 billion years ago, relates each of the 20 canonical amino acids to specific nucleotide triplets of genes and mRNAs and is an almost invariant (“frozen”) feature of life on Earth^6^. However, the genetic code likely rather started with a few small, simple amino acids that can be produced via a proto-metabolism, evolving gradually over time to add more complex amino acids to the repertoire with diverse functionalities. While the discovery of Selenocysteine and Pyrrolysine, the 21^st^ and 22^nd^ proteinogenic amino acids, implies some flexibility to the code, they are incorporated only at very specific positions in certain proteins. This begs the fundamental questions of why evolution stopped at these 20 amino acids^7–9^ and whether it is possible to alter the number and identity of these amino acids in a proteome-wide manner? Can a lineage with an altered set of canonical amino acids be evolved from an existing species? If we were able to experimentally alter natural species step by step so that they adopt different genetic codes with synthetic amino acids - then we would pave the way for the creation of artificial diversity and synthetic life.

Tryptophan is thought to be the latest addition to the genetic code with the highest metabolic cost of production^10,11^. Trp is one of only two canonical amino acids (cAAs) encoded by a single codon^12^. It is the least abundant amino acid in eukaryotic proteomes including humans (1.24% and 1.22%, respectively) and viruses (1.19%), and the second least abundant behind cysteine in both archean (1.03%) and bacterial proteomes (1.27%)^13^. By contrast, the most prevalent amino acid in all life forms, leucine, has an abundance between 8.84 % (viruses) and 10.09% (bacteria)^13^. Accordingly, Trp-residues have long been targets to demonstrate the chemical mutability of the genetic code by replacing them with noncanonical synthetic analogs^1,2^. In 2014, Yu et al.^14^ sequenced and analyzed a strain from an adaptation experiment of a Trp-auxotrophic *Bacillus subtilis* strain towards the noncanonical amino acid (ncAA) 4-fluoro-tryptophan (4-F-Trp) published more than 30 years earlier^2^. Only a few rounds of selection were necessary to give rise to a 4-F-Trp-tolerating strain with mutations of rpoC, in the sigma factors *sigI* ^15^ and *sigB* ^16^ as general stress sigma factors in *B. subtilis*. Most recently, a similar experiment performed with Trp-auxotrophic *Escherichia coli* revealed point mutations in the genes encoding non-essential vegetative sigma factors *rpoA* and *rpoC*, which help to reconfigure transcription under different environmental conditions, including stress^17^.

In this experimental vein, we have recently achieved a complete Trp substitution in the *Escherichia coli* proteome by its surrogate L-β-(thieno[3,2-*b*]pyrrolyl)-alanine ([3,2]Tpa, Figure 1) using adaptive laboratory evolution (ALE)^3^. Selective pressure was exerted on a Trp-auxotrophic strain which, in order to grow, was forced to perform conversion of [3,2]Tp and L-Ser to [3,2]Tpa through the reaction catalyzed by the endogenous enzyme Trp synthase (TrpS). However, when we tried to deduce the underlying adaptation mechanisms using ‘omics data, we realized that the genomic configuration of the ancestral strain we used was flawed. Specifically, it led to several additional selection pressures and subsequently led to off-target mutations that rendered a clear deduction of the genetic basis of the adaptation to [3,2]Tpa impossible (manuscript in prep).

**Figure 1.**
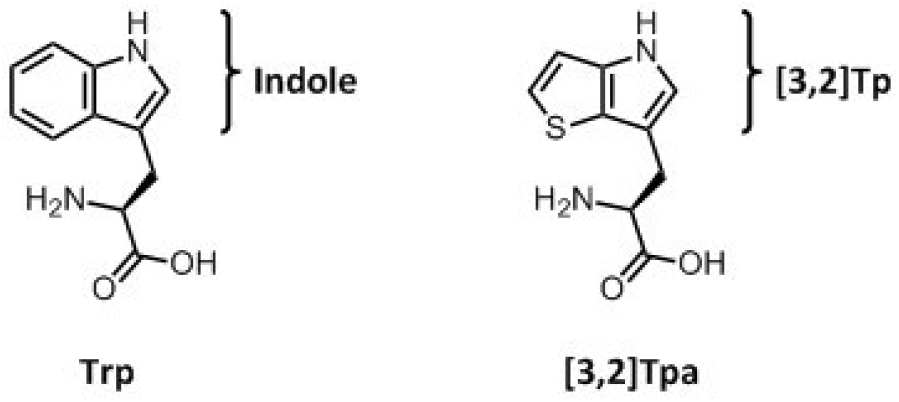
Chemical structures of Tryptophan and L-β-(thieno[3,2-*b*]pyrrolyl)alanine ([3,2]Tpa). The brackets indicate the structures of the precursors indole and β-thieno[3,2-*b*]pyrrole ([3,2]Tp).

Here, we have used what we learned from the first ALE experiment to design a novel ALE experiment using an *E. coli* strain with an optimized genomic configuration^18^. Using the new experimental framework, we were able to demonstrate that *E. coli* can succeed in overcoming the frozen state of the genetic code to replace tryptophan with the analog [3,2]Tpa, and identify that the primary adaptive mechanism is an alternation of the RpoS-mediated general stress response. The co-occurrence of mutations in stress regulators of Gram-positive bacteria supports the *E. coli* adaptation model towards Trp-analog usage presented here.

The general stress response is activated upon entering stationary phase and provides cross-protection to a wide spectrum of stresses *E. coli* are confronted with in their natural environments (*e*.*g*., nutrient limitation, competition of resources, fluctuating temperature, pH, osmolarity)^19–23^ The general stress response is mediated by the alternative sigma factor RpoS (σ^S^), which interacts with the core RNA polymerase (RNAP)^24^. RpoS governs the expression of a vast set of genes^25^ and shares control of various promoters with the vegetative sigma factor RpoD (σ^70^). The RpoS response is regulated at multiple levels that include transcription, translation, degradation, and activity^26^.

What happened during the evolution of bacteria and other microbes and organisms over several billions of years cannot be fully compared to the phenomenon of proteome-wide ncAA insertion experiments over the last 50 years^27^. Contemporary selection pressure of ncAA use and disposal is much more intense; selection is largely for survival in hostile minimal media environments rather than for traits providing fitness in slowly evolving populations in natural settings. In summary, what is occurring in our lifetime is an evolutionary process intensified by anthropogenic influences rather than the slower, random course of natural evolution.

During experimentally designed long-term serial dilutions, i.e. ALE, the exposure of bacterial metabolic prototypes to high concentrations of ncAAs for extended periods creates a severe selection pressure and leads to higher levels of incorporation. Knowledge of the intermediate steps in this important process would be revealing—how many steps are there from parent strain to fully adapted bacteria?

## RESULTS

### Adaptive Laboratory Evolution (ALE) on [3,2]Tp

We constructed an MG1655 derivative strain of *E. coli*, where the tryptophanase, as well as the majority of the Trp operon (*trpLEDC*), was knocked-out while retaining the tryptophan synthase (*trpBA*) on the chromosome under its natural regulation of *trpR* (MG1655 *ΔtnaA ΔtrpLEDC*). We refer to this new starting strain as TUB00 (Figure S1 A). The ALE was performed in new minimal medium (NMM)^28^.

In the first phase of the experiment, the concentration of indole was continually reduced; after only 29 passages (∼ 190 generations) the population was capable of growing in medium without the addition of any indole (Figure 2A). To avoid Trp contamination from commercial amino acid preparations, as described in other experiments^1^, in the second phase the other 19 canonical amino acids were removed from the medium in so-called metabolic blocks. After another 97 passages (∼ 630 generations) and a reduction of the cultivation temperature from 37°C to 30°C, methionine was the only canonical amino acid left in the medium. Within 10 additional serial dilutions and a total of 138 passages, the last canonical amino acid (cAA) was removed from the medium without loss of growth. During the last passages, the cultivation temperature was gradually increased back to 37°C (phase 3, Figure 2A). After a total of 170 passages, the adaptation experiment was terminated and the resulting strain was termed TUB170. We denote the media compositions in the following format throughout: NMM(a/b/c; a = number of cAAs; b = [3,2]Tp concentration in µM; c = indole concentration in µM). The ALE was initiated with NMM(19/25/1) and concluded with NMM(0/25/0) (Figure 2A).

**Figure 2.**
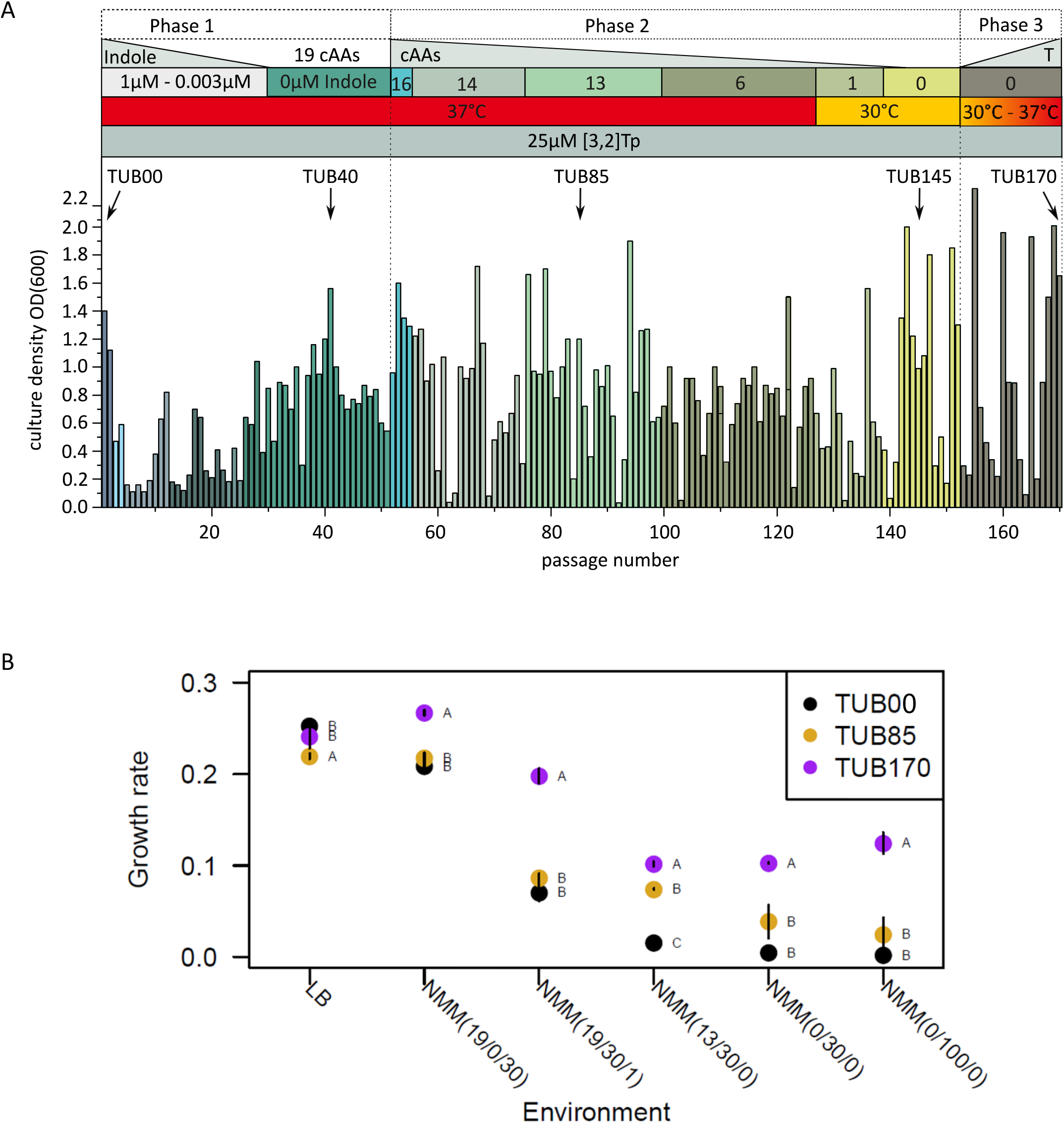
Adaptive Laboratory Evolution (ALE) on [3,2]Tp. A) Overview of optical densities throughout the adaptation experiment and the corresponding cultivation conditions. Phase 1: the indole concentration was reduced from 1µM to 0µM in several steps. Phase 2: All other 19 cAAs were removed according to metabolic blocks, whereby the temperature needed to be reduced to 30°C for removal of the last amino acid, Met. Phase 3: The temperature was gradually increased back to 37°C. B) Doubling times of TUB00, TUB85, and TUB170. Data were collected at 30 °C. The letters A, B, C indicate Tukey test results (statistical significance, p < 0.05), when different strains have different letters they are statistically different.

The growth rates of TUB00, TUB85 (isolated after 85 passages in phase 2), and TUB170 were compared in different media compositions (Figure 2B). TUB170 had a significantly higher growth rate than TUB00 and TUB85 in all NMM environments including the starting condition medium (NMM (19/30/1, Figure 2B). Interestingly, the growth rate of TUB170 increased when supplied with 100 µM [3,2]Tp (NMM(0/100/0)) compared to 30 µM [3,2]Tp (NMM(0/30/0)). A higher efficiency of the tRNA^Trp^ aminoacylation to [3,2]Tpa-tRNA^Trp^ or of [3,2]Tp conversion to [3,2]Tpa by the endogenous TrpBA could be a reason for this observation. Relative to TUB00 and TUB85, TUB170 had a significantly decreased growth rate in the standard rich medium LB (Figure 2B), indicative of a growth trade-off. TUB85 had the same growth rate as TUB00 except in the environment it was isolated in (NMM (13/30/0)), when it had an intermediate growth rate (Figure 2B), suggesting that this isolate contains a beneficial mutation that is specific to that environment.

### Mutations occurred in a wide range of genes

To characterize genetic changes that arose during the ALE, single colony isolates from different experimental phases were isolated and whole genome sequenced: TUB40 (phase 1), TUB85 (phase 2), TUB145 (phase 2), and TUB170 (phase 3). We identified 42 mutations within these four isolates, which provides a snapshot of the evolutionary dynamics of the ALE (Figure 3A, Table S4). An examination of the shared mutations (defined as any mutation that was present in at least two isolates) and private mutations (those present in only a single isolate) indicates that multiple lineages were likely present for the majority of the 170 transfers of the ALE (Figure 3B). Nine mutations were found in all sequenced isolates, indicating they arose in the first 40 transfers in the same genetic background (lineage) that spawned all sequenced isolates. These nine mutations occurred in genes involved in amino acid biosynthesis and processing (*aroG, astB*, and *leuS*), cell morphogenesis (*fliD, cvrA*), membrane proteins (*ylcJ, yejM*), oxidoreduction (*ydfI*), and in the promoter region of *yobF*, a small protein with unknown function.

**Figure 3.**
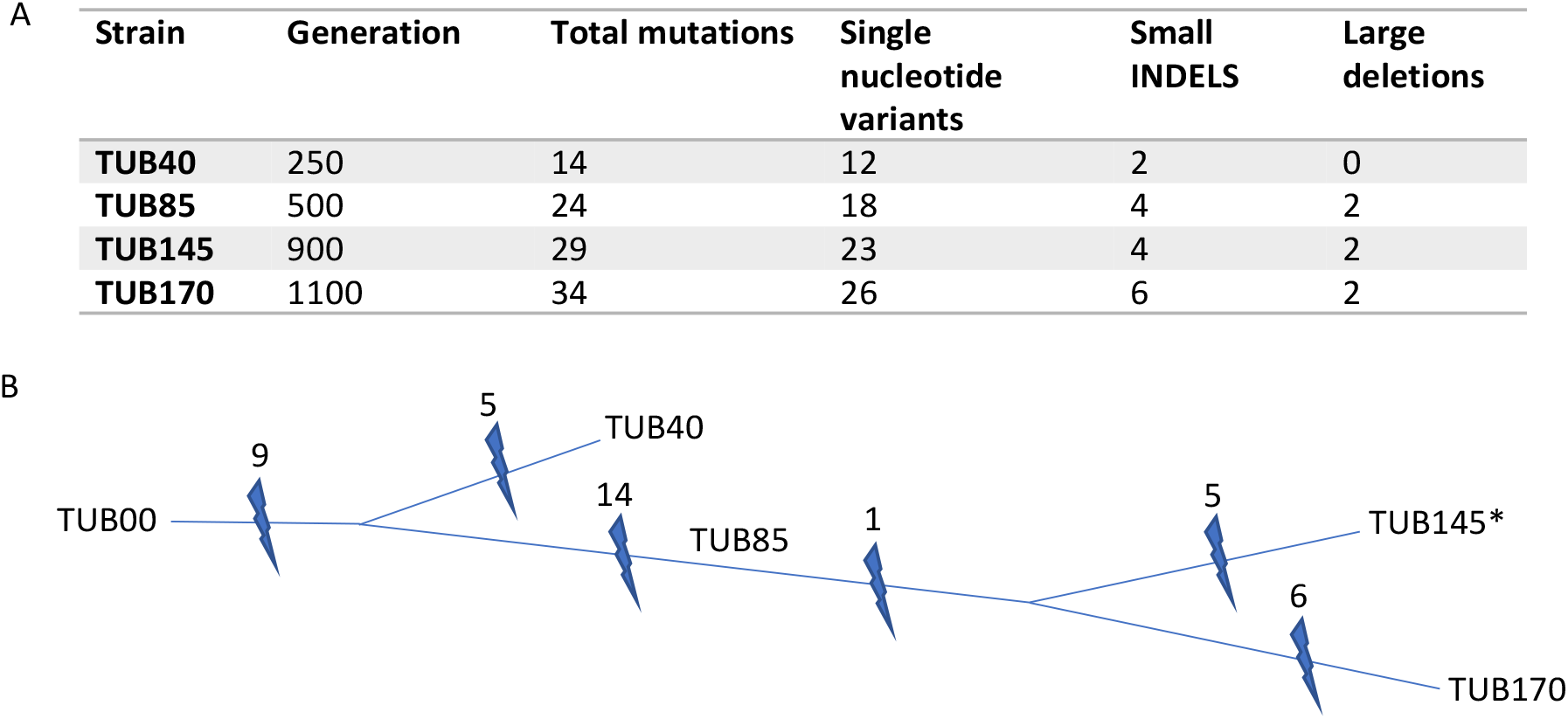
Genomic analysis of the ALE. A) The number of mutations identified in each sequenced isolate. Generations are estimated as described elsewhere^29^. B) A dendrogram depiction of the evolutionary history of the sequenced strains from the ALE experiment as they accumulate mutations (lightning bolt).

At some point before transfer 40, there was a split into at least two lineages. One led to TUB40, which has five private mutations. The second lineage was ancestral to the other three sequenced isolates (TUB85, TUB145, TUB170). The second lineage contains 13 shared single nucleotide variants in kinases (*glcK, pyrH, phoR*), phosphatases (*gpp*), reductases (*nfeF*), proteases (*lon, ftsH*), stress response factors (*rpoS, lrhA, gpp, sanA, sgrR*), and a mutation in a pseudogene (*ybfl*) and two large deletions. The larger is a 23 kb deletion involving 35 genes including the Rac prophage. The second is a 6 kb deletion involving 10 genes, beginning within the putative defective integrase of the Qin prophage (*intQ*) and ending within the putative selenite reductase (*ynfE*). No private mutations were identified in TUB85, indicating that this isolate is a common ancestor to both TUB145 and TUB170.

Only a single common mutation, in *ppc* (a phosphoenolpyruvate carboxylase), was found in both TUB145 and TUB170, suggesting that these two lineages split shortly after transfer 85. An additional four (TUB145) and nine (TUB170) private mutations were found in these two isolates. Interestingly, two genes with different mutations were found in these two terminal isolates: *cysN* (sulfate adenylyltransferase) and *gltA* (citrate synthetase), suggestive of strong selection for variants in these genes.

### Mutations involved in stress response

The mutation in the stress response gene *rpoS* (present in TUB85, TUB145, TUB170) particularly caught our attention. The mutation occurred at D118 (Figure 4A), which abolishes the salt bridge between D118 in σ^S^ and R275 in the RpoC-subunit of the core-RNAP (Figure 4B). The aspartate is conserved in most sigma factors (σ^S^, σ^D^, σ^H^) and is an important binding motif. To verify that an altered stress response is beneficial for the adaptation to [3,2]Tp, the *rpoS* gene was deleted in TUB00 via CRISPR/Cas9. We reasoned that reduced σ^S^-mediated stress response will improve the growth on the indole-analog compared to the wildtype. Therefore, the *ΔrpoS* strain, TUB170, and TUB00 were grown in media with varying indole concentrations. The optical density of the *ΔrpoS* strain was higher than that of the ancestral strain under all conditions (Figure 4 C), indicating increased tolerance towards the adverse effects of [3,2]Tp supplementation. Notably, even in media completely devoid of a Trp source (i.e., NMM(19/30/0) and NMM(0/30/0)), considerable growth of the *ΔrpoS* strain can be detected.

**Figure 4.**
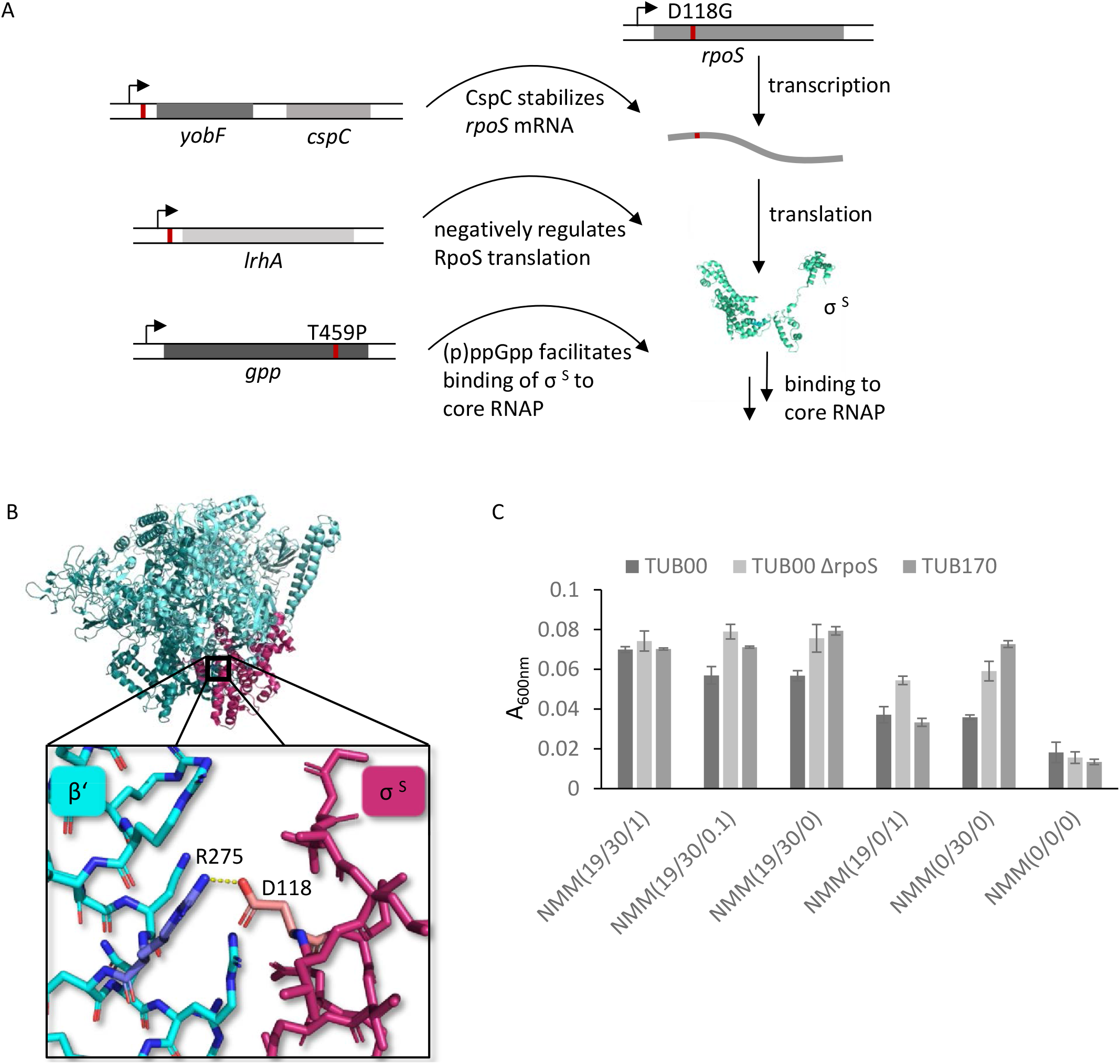
Mutations involved in the RpoS-mediated stress response. A) Schematic representation of the mutations directly involved in the general stress response and their impact on the stress response. B) The interaction between pseudo-Asp118 (rose) of the sigma factor S and Arg275 (blue) of the RpoC subunit of the core RNAP (PDB: 6OMF). The salt bridge is abolished by the mutation to Gly. C) Optical densities of the *ΔrpoS* mutant compared to the ancestral strain and the adapted strain TUB170 in the presence of varying indole concentrations after 48 h. NMM(19/30/1): starting conditions, NMM(19/30/0): end of phase 1, NMM(19/0/1): without [3,2]Tp, NMM(0/30/0): end conditions, NMM(0/0/0): without amino acids or precursors.

Additional general stress response proteins were also identified. Two of the mutations that arose before transfer 40 and were present in all sequenced isolates occurred in the promoter regions of the transcription units *yobF-cspC* and *lrhA*. The stress protein CspC stabilizes the *rpoS* mRNA^30^, while the DNA-binding transcriptional dual regulator LrhA represses *rpoS* translation^31^. A threonine to proline mutation in *gpp* was found in TUB85, TUB145, and TUB170, which the Protein Variation Effect Analyzer (PROVEAN)^32,33^ predicts to be deleterious. GppA hydrolyzes the 5’ phosphate from pppGpp to produce ppGpp, thereby fine-tuning the ratio between the two nucleotides^34^. This alarmone accumulates in response to amino acid (or other) starvation as part of the stringent response, which is also connected to the σ^S^-mediated general stress response^35^. The synthesis and stability of RpoS is enhanced by (p)ppGpp, thus boosting the expression of genes involved in stress protection^36,37^, and accumulation of (p)ppGpp or RpoS results in diminished fitness under nutrient-limited conditions^38^. Furthermore, (p)ppGpp increases the availability of core RNAP^39^, while indirectly decreasing cellular RpoD, thereby promoting binding of RpoS to RNAP^40,41^.

### Mutations deactivating proteases accompany the adaptation

Three mutations that arose in protease genes were predicted to be deleterious by PROVEAN (*lon* and *ftsH* were identified in TUB85, TUB145, and TUB170; *clpP* was found only in TUB170). We hypothesize that due to its physicochemical differences to Trp, the incorporation of [3,2]Tpa into proteins potentially yields locally or globally misfolded proteins, as reduction of protein stability upon [3,2]Tpa incorporation has been previously observed.^42^ Such misfolded proteins are usually cleared by proteases. However, in our novel strain, every single Trp position is replaced by [3,2]Tpa, presumably giving rise to a vast number of sub-optimally folded proteins. During the adaptation experiment, it could thus be advantageous for the bacterial system to lower this proteolytic activity in order to enhance the half-life of some affected proteins. The accumulation of mutations in proteases, particularly early in the ALE, implies that impaired protein folding is a concern when replacing Trp with [3,2]Tpa in the proteome and that downregulation of proteolytic activity might be beneficial for adaptation towards ncAA-usage.

### Proteomic analysis reveals major upregulation of stress proteins

Proteomic analysis was performed using stable isotope labeling by amino acids in cell culture (SILAC). Since SILAC relies on the labeling of proteins by the isotope-labeled amino acids ^13^C_6_^15^N_4_-arginine and ^13^C_6_^15^N_2_-lysine, the biosynthesis of arginine and lysine in the ancestral strain TUB00 was disabled via phage P1 transduction using strains from the Keio collection as donors^43^. The resulting strain was designated TUB00dKO and used as the reference in all SILAC experiments. By mixing the digested proteins from cells cultivated with and without the isotope-labeled amino acids, followed by quantification via high-resolution mass spectrometry, expression differences of proteins from cells cultured under different conditions can be analyzed.

To characterize the impact of [3,2]Tp addition on the proteome of the ancestral strain (TUB00), cells were cultivated in indole-rich medium without any [3,2]Tp (NMM(19/0/30)) and compared to the reference (TUB00dKO) cultured in medium supplied with [3,2]Tp mirroring the starting conditions, but also supplemented with the two isotope-labeled amino acids (NMM(17/30/1) + ^13^C_6_^15^N_4_-arginine and ^13^C_6_^15^N_2_-lysine; i.e., NMM(19/30/1) with two labeled amino acids).

Eighty-seven (87) proteins were up- or downregulated (log_2_-value > ± 2.5) in the absence of [3,2]Tp (Figure S2). Eight of the 36 upregulated proteins are related to chemotaxis (e.g., CheW, CheA) and motility (e.g., FliC, FliN, FliA, FliL; Table 1), a common response upon entering stationary phase and also a part of the general stress response^44–46^. The majority of the 51 most downregulated proteins are either directly connected to the general stress response under the control of the alternative sigma factor RpoS or via participation of the DNA-binding transcriptional dual regulator H-NS (Figure S2, cluster plot). H-NS is involved in fine-tuning RpoS expression under stress conditions, both by supporting RpoS-dependent expression^47–50^ over expression mediated by the housekeeping sigma factor RpoD^51–53^ and by destabilizing RpoS protein^48,54,55^. Downregulated proteins involved in the RpoS-mediated general stress response include proteins from the glutamate-dependent acid resistance system: GadC, GadB, GadA (Table 1). GadAB is a glutamate decarboxylase, which scavenges one proton upon decarboxylation of glutamate to γ-aminobutyrate (GABA)^56,57^, which is then secreted by the glutamate/GABA antiporter GadC.^56,58^ Another downregulated protein connected to the general stress response is the multidrug efflux transporter MdtEF (Table 1), involved in detoxification and described to be induced by indole.^59^

**Table 1.** Overview of up- and downregulated proteins in response to [3,2]Tp-supplementation. Isolates from the beginning of the ALE (TUB00, cultivated in NMM(19/0/30)) are compared to isolates from the middle (TUB85, cultivated in NMM(13/30/0)) and the end of the experiment (TUB170, cultivated in NMM(0/30/0)). The reference(TUB00dKO) was cultivated in a medium supplied with [3,2]Tp mirroring the starting conditions (NMM(19/30/1)). The change in abundance is reported as log2-fold and is visualized in the form of a heat map, where yellow indicates upregulation and blue indicates downregulation, white indicates no change in abundance.

| Function | Gene | Protein | Legend |  |  |
| --- | --- | --- | --- | --- | --- |
|  |  |  | upregulated |  | downregulated |
|  |  |  | Change of abundance in |  |  |
|  |  |  | the absence of [3,2]Tp TUB00 log2-fold | ALE isolate vs TUB00 TUB85 log2-fold | TUB170 log2-fold |
| chemotaxis | *cheW* | Chemotaxis protein CheW | -4.5 | n.a. | 0 |
|  | *cheA* | Chemotaxis protein CheA(L) | -3.7 | 0 | 0 |
| motility | *fliC* | Flagellar filament structural protein | -4.8 | 1 | 4.8 |
|  | *fliA* | RNA polymerase, sigma 28 (sigma F) factor | -3.4 | n.a. | 0 |
|  | *fliL* | Flagellar protein FliL | -1.9 | n.a. | n.a. |
|  | *fliG* | Flagellar motor switch protein | -3 | n.a. | 0 |
| acid resistance | *gadC* | L-glutamate:4-aminobutyrate antiporter | 5.3 | 6.1 | 6.5 |
|  | *gadB* | Glutamate decarboxylase B | 5.6 | 6.9 | 8.4 |
|  | *gadA* | Glutamate decarboxylase A | 3 | 5.9 | 0 |
| multidrug efflux transporter | *mdtE* | Multidrug efflux pump membrane fusion protein | 5.5 | 4 | 7 |
|  | *mdtF* | Multidrug efflux pump RND permease | 3.8 | 2.9 | 3.9 |
| Pho regulon | *phoU* | Phosphate signaling protein | -1.1 | -4.2 | -3 |
|  | *phoA* | Alkaline phosphatase | n.a. | -3.6 | -2.6 |
|  | *phoB* | DNA-binding transcriptional dual regulator PhoB | 0.7 | -2.7 | -2.8 |
|  | *pstS/phoS* | Phosphate ABC transporter periplasmic binding protein | 1.2 | -3.2 | -2.9 |
|  | *pstB* | Phosphate ABC transporter ATP binding subunit | -0.6 | -3.1 | -3.1 |
|  |  |  | upregulated |  | downregulated |
|  |  |  | Change of abundance in |  |  |
|  |  |  | the absence of [3,2]Tp TUB00 log2-fold | ALE isolate vs TUB00 TUB85 log2-fold | TUB170 log2-fold |
|  | *ugpB* | *sn*-glycerol 3-phosphate ABC transporter periplasmic binding protein | 0.2 | -2.5 | -3.6 |
| Leu biosynthesis | *leuA* | 2-isopropylmalate synthase | 0.2 | -4.6 | -3.1 |
|  | *leuD* | 3-isopropylmalate dehydratase subunit LeuD | -3.2 | -4.5 | -3.1 |
|  | *leuB* | 3-isopropylmalate dehydrogenase | 0.1 | -4.5 | -1.7 |
|  | *leuC* | 3-isopropylmalate dehydratase subunit LeuC | -3.5 | -4 | -2.4 |
| Trp synthesis | *trpA* | Tryptophan synthase subunit $\alpha$ | -2.3 | -2.2 | -1.6 |
| | *trpB* | Tryptophan synthase, $\beta$ subunit dimer | -1.7 | -1.9 | -1.6 |
| RpoS repressor | *lrhA* | DNA-binding transcriptional dual regulator LrhA | -1.4 | -2.5 | -1.1 |
| *rpoS* stabilizer | *cspC* | CspA family stress protein CspC | -1 | 3.9 | 3.7 |
| galactose catabolism | *galE* | UDP-glucose 4-epimerase | 5.7 | 5.3 | 5 |
|  | *galK* | Galactokinase | 5.2 | 4.9 | 4.7 |
|  | *galT* | Galactose-1-phosphate uridylyltransferase | 4.6 | 4.8 | 4.8 |
| protection oxid. and osmotic stress | *osmC* | Peroxiredoxin OsmC | 4.6 | 3 | 3.2 |
| pyruvate metabolism | *poxB* | Pyruvate oxidase | 4 | 6 | 3.9 |

To investigate how the expression profile changed over the course of the adaptation experiment we cultivated the adapted strains TUB85 and TUB170 in media analogous to the starting conditions (NMM(19/30/1)), as well as in their adapted media (NMM(13/30/0) and NMM(0/30/0), respectively) to distinguish between effects caused by the removal of the cAAs from the media and those responsible for adaptation to [3,2]Tp. TUB85 and TUB170 samples yielded similar expression profiles under all three culturing conditions, with the only major difference being an expected upregulation of amino acid biosynthesis enzymes in media lacking cAA supplementation (Figures S3 and S4). Thus, while the ancestral strain reacted to the presence of [3,2]Tp by inducing the RpoS-mediated general stress response, both adapted strains seem to re-regulate this response and restore an unstressed ground state similar to the one observed for the ancestral strain in the medium lacking [3,2]Tp (Table 1). Compared to the ancestral strain cultivated in [3,2]Tp-supplemented media, 105 proteins are up- or downregulated in TUB85 and TUB170 in the presence of [3,2]Tp. Of these, 39 proteins are upregulated (Figure S3) and comprise proteins under the regulation of the Pho-regulon (PhoU, PhoA, PstS, PhoB, PhoS, PstB, UgpB), which are usually responsible for P_i_ uptake in phosphate limiting environments^60^. As the cultivation media used in this study contain phosphate buffer in non-limiting concentrations, it is not clear why these proteins are upregulated. We hypothesize this could be a response to the mutation in *sgrR* which activates the small RNA gene *sgrS* under glucose-phosphate stress conditions. Induction of the Pho Regulon has been shown to suppress the growth defect of an *Escherichia coli sgrS* mutant^61^. Others have shown a link between the Pho regulon and the synthesis of polyphosphate and guanosine tetraphosphate (ppGpp), a key phosphate-containing molecule involved in the stringent response, which is responsible for the inhibition of RNA synthesis when there is a deficiency in the amino acid availability^62^. The Pho regulon is repressed in *E. coli* mutants that cannot accumulate ppGpp although the actual mechanism relating these two is unclear^63^.

Proteins from the leucine biosynthesis pathway are also upregulated in TUB85 and TUB170. This pathway is activated by the transcriptional regulator LeuO^64^, which also indirectly inhibits *rpoS* translation by repressing the transcription of the sRNA DsrA.^65^ DsrA, in turn, stabilizes *rpoS* mRNA^66^ and also activates its translation^67–69^. Perhaps not surprisingly, another prominent protein upregulation in the adapted strains is that of the tryptophan synthase TrpBA, which is responsible for the Trp synthesis from indole and serine^70^ (or in this case [3,2]Tpa synthesis from the precursor [3,2]Tp and serine^3,42^). Furthermore, LrhA, for which a mutation in the promoter region was detected, is also upregulated. LrhA is a repressor of *rpoS* translation^31^.

The most prevalent feature among the 66 downregulated proteins revolves around the σ^S^-mediated general stress response (Figure S4), thus reversing the effect [3,2]Tp elicits at the beginning of the adaptation experiment. One such example is the downregulation of the *rpoS* mRNA-stabilizing protein CspC, which is accompanied by a mutation in its promoter region (Figure 4 A).

## DISCUSSION

Here we set out to ask whether we could evolve a lineage with an altered set of canonical amino acids from an existing species, a feat that could pave the way for the creation of artificial diversity and synthetic life. To accomplish this, we performed an ALE with an *E. coli* MG1655 derivative to obtain cells with chemically evolved proteomes in which one amino acid has been completely replaced. We chose tryptophan (Trp) as our canonical target for substitution by the physicochemically similar, non-canonical, analog L-β-(thieno[3,2-*b*]pyrrolyl)alanine ([3,2]Tpa)^3^. Since Trp is rarely an essential catalytic moiety in *E. coli* enzymes, it is well suited for such experiments. The initial medium was NMM supplemented with indole and amino acids. After 170 transfers, our terminal evolved strain was able to grow in NMM + 25 µM [3,2]Tp without additional supplements.

In the initial phase of the experiment (before transfer 40, generation 250) the concentration of indole was reduced from 1 µM to zero. *E. coli* likely adapted to both the presence of the non-canonical amino acid precursor [3,2]Tp as well as concomitant Trp starvation. We observed three mutations in genes with a direct link to amino acid metabolism (*astB, aroG*, and *leuS*). The first mutation introduced a stop codon leading to a truncation of the N-succinylarginine dihydrolase (AstB). *astB* is a member of the ammonia-producing arginine succinyltransferase (*astCADBE*) operon which is necessary for arginine degradation during nitrogen-limited growth and also contributes to the degradation of other amino acids. As the growth medium used in this study includes a non-limiting ammonia source (Ammonium sulfate), it is essential to reduce the ammonia producing capacity of this operon while maintaining its ability to perform its other functions. Disruption of *astB* has been shown to eliminate succinylarginine dihydrolase activity and prevent arginine utilization without impairing ornithine catabolism^71^. This reduced dependency on arginine as a nitrogen source enables arginine to serve other relevant functions such as the biosynthesis of the most common polyamines, putrescine and spermidine, which are required for optimal growth through their involvement in several physiological processes^72^.

In the second phase of the experiment, the number of canonical amino acids was decreased from 19 to 0. Between transfer 40 (generation 250) and transfer 85 (generation 500), the evolving population acquired many mutations in genes related to chemotaxis and flagella synthesis. It is typical for *E. coli* to cease synthesis of chemotaxis and flagella upon entering stationary phase or being exposed to stresses^44–46^ while increasing the production of stress-related proteins and drug exporters. In this state, *E. coli* redirects its metabolism from producing primarily growth-related (macro)molecules to using its limited resources for maintenance and survival^23^.

In this second phase, the evolved population also acquired a mutation in *rpoS* that lead to RpoS inactivation contributing to improved adaptability. In particular, mutation of the RpoS Asp118 residue abolished RpoS’ capacity for repression, diminishing the RpoS-mediated general stress response. *rpoS* mutations observed in a laboratory context should be evaluated critically, as storage and shipment of strains in stab cultures or glycerolized cultures on filter disks are known to select for *rpoS* mutants^73,74^. Nevertheless, the *rpoS* mutation and concomitant downregulation of RpoS protein detected in this study are unlikely to be such storage-related artifacts. Samples taken from the ALE experiment were immediately frozen at -80°C and isolates used for later characterization and analysis were directly derived from these frozen fossils. Indeed, deletion of *rpoS* in the ancestral strain is sufficient for the growth of the un-adapted strain on [3,2]Tp, and *rpoS* mutations have been identified in previous ALE experiments in nutrient-limited conditions, such as in glucose-limited chemostats.^75^ A similar experiment was conducted recently with a genome-reduced MG1655 derivative showing growth defects in minimal medium. The deletion of *rpoS* restored the growth rate of the unevolved strain to 80% of the growth rate of the evolved strain^76^. High amounts of RpoS protein typical in stationary phase cells or cells under stress cause competition between RpoS and the vegetative sigma factor RpoD for a limited number of RNA polymerase core subunits. Mutations in *rpoS* therefore likely alleviate the competition in favor of nutrient scavenging through increased expression of RpoD-dependent genes.

In a natural habitat prone to fluctuations in environmental conditions such as temperature, osmolarity, or pH, loss-of-function of RpoS is likely to be disadvantageous. Loss of function of RpoS can also be detrimental under certain stress conditions^77^. For example, *rpoS* mutants evolved under osmotic stress insert IS*10* in the promoter of the *otsBA* operon, thereby partially restoring the wildtype response to osmotic stress by altering the normally RpoS-dependent expression to a RpoS-independent one^78^. In other instances, the appearance of attenuated *rpoS* mutations in long-term stationary phase cultures^79^, as well as in glucose-limited chemostats under acidic pH^75^ suggests less drastic mutations may be beneficial. These findings, together with the fact that *rpoS* deletions seem to be rare in natural *E. coli* populations^80^, indicate that the nature of the trade-off between reduced stress resistance and increased nutrient scavenging is part of a very careful equilibrium.

In the isolates TUB85 and TUB170, we observed the upregulation of proteins from the leucine biosynthesis pathway (*leuLABCD* operon). Transcription of this operon is not only activated by LeuO but also guanosine tetraphosphate (ppGpp) which activates the binding of RNA polymerase to the *leuLABCD* promoter (*leuLp*) under conditions of leucine starvation. Interestingly, the leucine supplement was suspended for these two strains. ppGpp is produced by guanosine-5’-triphosphate,3’-diphosphate phosphatase (GppA) and its gene acquired a T459P mutation in our ALE experiment. Thus, the upregulation of proteins from the leucine biosynthesis pathway could be the hyperactivation of the *leuLABCD* operon transcription by both LeuO and ppGpp. Additionally, aside from activating leucine biosynthesis, LeuO indirectly inhibits *rpoS* translation, thereby deactivating the general stress response.

*E. coli* mitigated its stress response in favor of cell growth when adapted to a progressive depletion of an essential canonical substrate. The proteomic data revealed that the levels of stress-related proteins subside to levels corresponding to the ancestral strain in relaxing media during the time-course of the adaptation experiment. This discovery is supported by genomic data identifying mutations related to four stress proteins, including two up- or downregulated proteins and the master regulator RpoS itself. In order to neglect the negative effects of [3,2]Tpa incorporation, the adapted strain lowers its protein quality management by reducing its proteolytic activity as evinced by mutations in three proteases.

Our study shows that adaptation to proteome-wide ncAA-insertions is highly pleiotropic. Pleiotropic interactions can result from changes in the metabolic pathways that are networked in the cells; e.g. a change in the concentration of an enzyme or protein could lead to the reconfiguration of intracellular networking. These interactions are an integral part of the adaptation mechanisms associated with proteome-wide insertions and provide the means to attenuate stress responses, modulate folding pathways or other functions. Translational activity is only one of the many biological functions of canonical amino acids; ncAA integration will influence not only proteomes but also cell physiology, signaling, and metabolic and structural processes.

Our findings contribute a step towards understanding possible environmental causes of genetic changes and their relationship to evolution. The knowledge of how cellular metabolism can be regulated in response to alterations of the environment uncovered in this study might help guide engineering efforts and the creation of synthetic cells.

## ACKNOWLEDGMENTS

I. T. and S. O. were supported by the GRK 1582 “Fluorine as a Key Element”, funded by Deutsche Forschungsgemeinschaft (DFG). N. B. thanks Christian Thomsen, President of the Technical University of Berlin, and Canada Research Chairs for support (grant no. 950-231971). A. C. G. and A-R. A. B. were funded by an NSERC Discovery Grant to A. C. G. L. P. was supported by a Marie Curie Intra European Fellowship within the 7th European Community Framework Programme and by an Estonian Research Council grant (PUT626). The Wellcome Centre for Cell Biology is supported by core funding from the Wellcome Trust [203149].

## AUTHOR CONTRIBUTIONS

I. T. wrote the final version of the manuscript and provided samples for additional and control experiments. M. H., N. B., and S. O. designed the experimental setup, and S. O. performed the serial dilution experiments and associated analytical/microbiological procedures. A. G., M. H., and S. O. contributed to the writing of the manuscript at various stages. A. G., T. S., and A-R. A. B. performed genetic analyses and constructed pedigrees. L. P. and J. R. performed proteomic analyses, and together with S. O. interpreted the data. C. T.-K. performed chromosomal gene deletions and the corresponding growth assays. N. B. conceived the project, directed the experimental work, and wrote the first version of the manuscript.

## DATA AND MATERIALS AVAILABILITY

Materials and methods, as well as additional data, are available in the separate SI file.

