## Supplementary material for "Evolving a mitigation of the stress response pathway to change the basic chemistry of life": SI

### Spplementary Tables

### 1. MATERIALS AND METHODS

#### Inoculation method of the adaptive evolution experiment

In a 100mL Erlenmeyer flask covered with aluminum foil 10mL of the respective NMM-medium were inoculated 1:100 with the previous culture. The culture was grown under agitation for 1 or 2 days until the $OD_{600}$ was constant. Medium supplements with indole and amino acids were removed as fast as possible while keeping the population density above 0.1 $OD_{600}$ to avoid bottleneck effects.

#### Doubling times of TUB-strains

Isolates were re-grown from a cryostock in 5mL LB at 37°C overnight. To remove the LB medium, the cells were washed twice with NMM0. 10mL of the respective NMM were inoculated with 100 µL of these NMM0 washed cells and grown at 30°C or 37°C until stationary phase was reached (generally between 1-2 days). This step was repeated twice and subsequently, these cultures were used to inoculate a 96-well plate with 200 µL cultures in a 1:100 ratio. $OD_{600}$ values were obtained at the Infinite®M200 plate reader (Tecan Group AG, Männedorf, Switzerland). The maximum growth rate was calculated from a custom spline-fitting R script. Please note that while during the adaptation experiment a [3,2]Tp concentration of 25 µM was used, a slightly higher indole analog concentration of 30 µM was used for the characterization experiments due to increased culturing stability.

#### Next generation sequencing and analyses

Genomic DNA was sent to Beijing Genomics Institute in Hong Kong for re-sequencing. For TUB40, TUB85, and TUB145 an Illumina HiSeq 2000 and for TUB170 an Illumina HiSeq 2500 sequencing platform was used. The sequencing data were aligned to E. coli MG1655 (NC_000913.3) and evaluated by the company.

Evidence of evolution of the strains was assessed using breseq v0.34.0 (Deatherage and Barrick, 2014) running on 15 processors in consensus mode with a limit-fold-coverage (-l) of 100. Consensus mode is appropriate when re-sequencing a clonal haploid genome. MG1655 (NC_000913.3) was used as the reference in all analyses. The number of mutations in each genome was counted using the gdtools utility of breseq. The gdtools utility was also used to consolidate all the mutations from the individual strains into one gd file which was then converted into a vcf file and the effect of mutations was evaluated with SnpEff[1].

The phylogeny of the clones was constructed with MG1655 as the outgroup. Briefly, gdtools COMPARE was used to generate a multiple sequence alignment phylip file from the gd file outputs from breseq analyses of the strains. The phylip file was imported into Seaview (Gouy et al., 2010) and the alignment statistics were obtained. A phyML tree was then constructed with a GTR model, 100 boostraps, with the best of NNI and SPR as tree searching operation. The tree was then visualized with iTOL[2].

**Cluster plots using string.db**

Cluster plots were generated using the STRING v10 database (http://string-db.org/). This bioinformatics tool allows seeing interactions of a pair of proteins based on data mining. Please find a detailed description of the software in Szklarczyk et al.[3]

**Chromosomal gene deletions: Phage P1 transduction and CRISPR/Cas9**

5mL LB were inoculated 1:100 with an overnight culture of the donor strain and the cells were grown under agitation at 37 °C to an $OD_{600}$ of around 0.6. After addition of 5mM $CaCl_2$ and incubation for 30 min at 37 °C, 100 µL of the cell suspension were mixed with 100 µL of a solution of phage P1. After incubation at 37 °C for 20 min without shaking, the mixture was added to 4 mL 0.6 % soft agar supplemented with 5mM $CaCl_2$. The soft agar-containing infected donor cells was poured on LB agar plates and incubated overnight. After the soft agar-containing plaques were removed, 800 µL $CHCl_3$ were added. The suspension was mixed vigorously. After centrifugation (15000 x g, 10 min), the supernatant was conserved by adding a few drops of $CHCl_3$ and was stored at 4 °C. The recipient strain was grown under agitation at 37 °C to an $OD_{600}$ of around 0.6. After addition of 5mM $CaCl_2$ and incubation for 30 min at 37 °C, 1mL of the cell suspension was infected with 30 µL of the donor phage solution. After incubation at 37°C for 15 min without shaking, the cells were pelleted and resuspended in 1mL LB supplemented with 0.1M sodium citrate. After shaking for 45 min at 37°C the cells were plated on $LB^{Kan}$ agar plates and incubated overnight at 37°C. The knockout was verified by colony PCR.

CRISPR/Cas9 deletions were conducted as previously described using the CAGO technique[6].

**Proteomic analysis - Sample preparation and measurement**

Isolates were re-grown from a cryostock in 5 mL LB- medium at 37 °C overnight. To remove residual LB-medium, the cells were washed twice with NMM0. 10 mL of the respective NMM were inoculated with 100 µL of these NMM0 washed cells and grown at 30 or 37 °C until stationary phase was reached (generally between 1-2 days). This step was repeated twice. Subsequently, those cultures were used to inoculate triplicates of 50 mL cultures. These cultures were grown to early stationary phase and cells were harvested by centrifugation (4 °C, 5.000 g, 15 min). Cell pellets were lysed in 100 mM TRIS (pH 7.5), 4% sodium dodecyl sulfate (SDS), and 100mM DTT at a temperature of 95 °C for 10 min, followed by sonication in a Bioruptor Standard (Diagenode, Belgium) for 30 sec on, 30 sec off, 10 cycles, power setting at high (H); samples kept on ice at all times. Lysates were clarified by centrifugation (4 °C, 17.000 g, 5 min).

'Heavy'-labeled (H) TUB00dKO (NMM(17/30/1) + $^{13}C_6{}^{15}N_4$-arginine and $^{13}C_6{}^{15}N_2$-lysine [Silantes, Munich, Germany]) lysate was mixed at 1:1 ratio with different 'light'-labeled (L) lysates - TUB00 (NMM(19/0/30)), TUB85 (NMM(19/30/1)), TUB85 (NMM(19/30/0)), TUB170 (NMM(19/30/1)), TUB170 (NMM(19/30/0)), proteins were digested as described[4].

For LC-MS/MS analysis, peptides were desalted on C18-StageTips[5] and separated by reversed-phase liquid chromatography on a Dionex UltiMate RSLCnano 3000 system coupled to Q Exactive mass-spectrometer (Thermofisher Scientific, USA). Peptides were loaded onto a 75 µm × 300 mm fused silica emitter (New Objective, USA) packed in-house with Reprosil-Pur C18-AQ 3 µm particles (Dr. Maisch, Germany) at a flow-rate of 500 nL/min for 10 minutes and separated using 230 min 2-40% B gradient (A: 0.1% formic acid, B: 0.1% formic acid/80% acetonitrile) at a flow-rate of 200 nL/min. Separating gradient was followed by an 11-min ramp to 95% B, wash at 95% B for 5 minutes, and return to initial conditions for the next run.

Eluted peptides were sprayed directly into a Q Exactive mass spectrometer operated in data-dependent mode with up to ten MS/MS scans (NCE=25) being recorded for each precursor ion scan. Precursor ion spectra were recorded in profile (m/z 350-1400, R = 70 000); data-dependent MS/MS spectra were acquired in profile (NCE 25, R = 17500). Mono-isotopic precursor selection was enabled, singly charged ions and ions with an unassigned charge state were rejected, each fragmented ion was dynamically excluded for 90 s.

MS data were analyzed with MaxQuant 1.3.0.5, using *Escherichia coli* MG1655 UniProtKB reference proteome sequence database (downloaded 20110909) with the following parameters: enzyme – Trypsin/P, precursor mass accuracy – 10 ppm, fragment mass accuracy – 20 ppm, fixed modification - Carbamidomethyl (C), variable modifications - Oxidation (M), Trp->[3,2]Tpa (W + 5.956 Da), FDR values were set at 0.01.

### 2. SUPPLEMENTARY TABLES

**Table S1 I Strains used throughout this study**

| | |
|---|---|
| **MG1655** | *E. coli* K12 F$^-$ λ$^-$ *ilvG$^-$ rfb$^-$*50 *rph-1* |
| **TUB00** | MG1655 Δ*trpLEDC::FRT* Δ*tnaA::FRT* |
| **TUB00-dKO** | TUB00 Δ*lysA::FRT* Δ*argA::FRT* |
| **JW2806-1** | E. coli F$^-$ Δ(*araD-araB*)*567*<br>*ΔlacZ4787(::rrnB-3)* λ$^-$ Δ*lysA763::kan*<br>rph-1 Δ(*rhaD-rhaB)568 hsdR514* |
| **JW2786-1** | E. coli F$^-$ Δ(*araD-araB*)*567*<br>*ΔlacZ4787(::rrnB-3)* λ$^-$ Δ*argA763::kan*<br>rph-1 Δ(*rhaD-rhaB)568 hsdR514* |

**Table S2 I ANOVA results for each environment.** Tukey test results indicate statistical significance ($p < 0.05$), when different strains have different letters they are statistically different.

| | | **Tukey Test Results** | | |
|---|---|---|---|---|
| **Environment** | **ANOVA model** | **TUB0** | **TUB85** | **TUB170** |
| LB | $F_{2,6} = 6.04$, $p = 0.037$ | B | A | B |
| 19-0-30 | $F_{2,6} = 85.62$, $p < 0.0001$ | B | B | A |
| 19-30-1 | $F_{2,6} = 112.2$, $p < 0.0001$ | B | B | A |
| 13-30-0 | $F_{2,6} = 1037$, $p < 0.0001$ | C | B | A |
| 0-30-0 | $F_{2,6} = 32.66$, $p = 0.0006$ | B | B | A |
| 0-100-0 | $F_{2,6} = 40.11$, $p = 0.0003$ | B | B | A |

**Table S3 I Composition of the different NMM media throughout the ALE.**

| Medium | c (Ind) [µM] | c ([3,2]Tp) [µM] | Supplemented amino acids | Passage |
|---|---|---|---|---|
| NMM19 | 1 | 25 | Ala, Arg, Asn, Asp, Cys, Gln, Glu, Gly, His, Ile, Leu, Lys, Met, Phe, Pro, Ser, Thr, Tyr, Val | 1-2 |
| NMM19 | 0.5 | 25 | Ala, Arg, Asn, Asp, Cys, Gln, Glu, Gly, His, Ile, Leu, Lys, Met, Phe, Pro, Ser, Thr, Tyr, Val | 3-4 |
| NMM19 | 0.2 | 25 | Ala, Arg, Asn, Asp, Cys, Gln, Glu, Gly, His, Ile, Leu, Lys, Met, Phe, Pro, Ser, Thr, Tyr, Val | 5-12 |
| NMM19 | 0.1 | 25 | Ala, Arg, Asn, Asp, Cys, Gln, Glu, Gly, His, Ile, Leu, Lys, Met, Phe, Pro, Ser, Thr, Tyr, Val | 13-21 |
| NMM19 | 0.05 | 25 | Ala, Arg, Asn, Asp, Cys, Gln, Glu, Gly, His, Ile, Leu, Lys, Met, Phe, Pro, Ser, Thr, Tyr, Val | 22-24 |
| NMM19 | 0.025 | 25 | Ala, Arg, Asn, Asp, Cys, Gln, Glu, Gly, His, Ile, Leu, Lys, Met, Phe, Pro, Ser, Thr, Tyr, Val | 25-28 |
| NMM19 | 0 | 25 | Ala, Arg, Asn, Asp, Cys, Gln, Glu, Gly, His, Ile, Leu, Lys, Met, Phe, Pro, Ser, Thr, Tyr, Val | 29-51 |
| NMM16 | 0 | 25 | Ala, Arg, Asn, Asp, Gln, Glu, His, Ile, Leu, Lys, Met, Phe, Pro, Thr, Tyr, Val | 52-55 |
| NMM14 | 0 | 25 | Ala, Arg, Asn, Asp, Gln, Glu, His, Ile, Leu, Lys, Met, Pro, Thr, Val | 56-74 |
| NMM13 | 0 | 25 | Ala, Arg, Asn, Asp, Gln, Glu, Ile, Leu, Lys, Met, Pro, Thr, Val | 75-98 |
| NMM6 | 0 | 25 | Arg, Ile, Leu, Met, Thr, Val | 99-126 |
| NMM1 | 0 | 25 | Met | 127-137 |
| NMM0 | 0 | 25 | | 138-170 |

**Table S4 I Identified mutations**

| TUB40 | TUB85 | TUB145 | TUB170 | Gene | Position | Mutation | Function |
|---|---|---|---|---|---|---|---|
| X | X | X | X | ylcJ | 568798 | Y35C (A→G) | Small protein expressed during exponential phase in rich growth medium |
| X | X | X | X | leuS | 674338 | V149A (T→C ) | Aminoacyl tRNA synthetase |
| X | X | X | X | aroG | 786075 | I148T (T→C) | Chorismate pathway, which leads to the biosynthesis of aromatic amino acids. |
| X | X | X | X | cvrA | 1240910 | Δ1 bp | K+ efflux, may function as a K+:H+ antiporter with a role in adaptive stress relief |
| X | X | X | X | ydfI | 1630102 | Δ1 bp | Putative oxidoreductase |
| X | X | X | X | astB | 1827488 | T258* (C→T) | Catalyzes the second reaction in the ammonia-producing arginine catabolic pathway |
| X | X | X | X | Intergenic (yobF/yebO) | 1907629 | -38/+632 (T→C) | DUF2527 domain-containing protein/uncharacterized protein |
| X | X | X | X | fliD | 2004929 | (G352D) G→A | Morphogenesis and elongation of the flagellar filament |
| X | X | X | X | yejM | 2285853 | D493V (A→T ) | Inner membrane protein implicated in lipid homeostasis and cardiolipin trafficking to the outer membrane. |
| x | | | | cra | 88887 | L287P (T→C) | DNA-binding transcriptional dual regulator Cra |
| x | | | | intergenic (thpR/hrpB) | 162044 | -13/-61 (G→C) | RNA 2',3'-cyclic phosphodiesterase/putative ATP-dependent RNA helicase |
| x | | | | sufB | 1762881 | I377V (A→G) | Fe-S cluster scaffold complex subunit |
| x | | | | rfaD | 3794366 | F127S (T→C) | ADP-L-glycero-D-mannoheptose 6-epimerase |
| x | | | | serB | 4625753 | I287V (A→G) | Phosphoserine phosphatase |
| | X | X | X | sgrR | 75651 | D550V (A→T) | Activates the small RNA gene sgrS under glucose-phosphate stress conditions as well as yfdZ. |
| | X | X | X | pyrH | 192068 | N72H (A→C)) | UMP kinase is an essential enzyme involved in the de novo biosynthesis of pyrimidine nucleotides. |

| TUB40 | TUB85 | TUB145 | TUB170 | Gene | Position | Mutation | Function |
|---|---|---|---|---|---|---|---|
| | X | X | X | phoR | 418083 | Δ1 bp | Indirectly senses and responds to variations in the level of extracellular inorganic phosphate |
| | X | X | X | lon | 459996 | A370V (C →T) | ATP-dependent protease, degradation of misfolded proteins as well as a number of rapidly degraded regulatory proteins. |
| | X | X | X | intergenic (*ybeF*/*lipB*) | 661,522 | T→C | Putative LysR-type DNA-binding transcriptional regulator YbeF/lipoyl(octanoyl) transferase |
| | X | X | X | ybfI | 737025 | T→C | putative transposase |
| | X | X | X | Δ23,060 bp | 1411925 | Δ23,060 b p | |
| | X | X | X | Δ6,805 bp | 1652172 | Δ6,805 bp | |
| | X | X | X | glxK | 1907428 | P101S (C→T) | Catalyzes the phosphorylation of D-glycerate |
| | X | X | X | sanA | 2233281 | V135A (T→C) | Participates in the barrier function of the cell envelope. |
| | X | X | X | Intergenic (lrhA/alaA) | 2406689 | -48/-872( A→G) | DNA-binding transcriptional dual regulator/glutamate--pyruvate aminotransferase |
| | X | X | X | rpoS | 2867199 | D118G (A→G) | A subunit of RNA polymerase that acts as the master regulator of the general stress response |
| | X | X | X | nfeF | 3215760 | R244R (G→A) | NADPH-dependent ferric reductase |
| | X | X | X | ftsH | 3326088 | D28G (A→G) | Part of the FtsH/HflKC complex, degrades both soluble and inner membrane proteins. |
| | X | X | X | gpp | 3962855 | T459 (A→C) | Catalyzes the conversion of pppGpp to ppGpp. |
| | | X | X | ppc | 4151253 | M616V (A→G) | Phosphoenolpyruvate carboxylase |
| | | x | | gltA | 754054 | T139A (A→G) | Citrate synthase |
| | | x | | cysN | 2875343 | S26N (G→A) | Sulfate adenylyltransferase subunit 1 |
| | | x | | rho | 3966607 | F64S (T→C) | Transcription termination factor |

| TUB40 | TUB85 | TUB145 | TUB170 | Gene | Position | Mutation | Function |
|---|---|---|---|---|---|---|---|
| | | x | | intergenic (yjdN/yjdM) | 4326088 | -347/311 (T→G) | Conserved protein/conserved protein |
| | | | x | clpP | 456933 | A86V (C→T) | ATP-dependent Clp protease proteolytic subunit |
| | | | x | fes | 613377 | V214A (T→C) | Enterochelin esterase |
| | | | x | gltA | 753958 | T171P (A→C) | Citrate synthase |
| | | | x | nrdB | 2347643 | I87T (T→C) | Ribonucleoside-diphosphate reductase 1 subunit beta |
| | | | x | bcp | 2600729 | Δ1 bp | Thiol peroxidase |
| | | | x | cysN | 2875127 | A98G (C→G) | Sulfate adenylyltransferase subunit 1 |
| | | | x | ubiH | 3053250 | V90A (T→C) | 2-octaprenyl-6-methoxyphenol hydroxylase |
| | | | x | cyaA | 3991157 | 2 bp→CT | Adenylate cyclase |
| | | | x | intergenic (nanC/fimB) | 4540077 | -576/-880 (A→G) | N-acetylneuraminic acid outer membrane channel/regulator for fimA |

### 3. SUPPLEMENTARY FIGURES

A

**Verification of ΔtrpLECD knockout in TUB00**

| | |
|---|---|
| 217 | *CTCCCGTTGCGTTGCATTG* |
| 218 | *CGCCGTGCAGCAAATCTTC* |
| 220 | *CACTGCATAATTCGTGTCGCTC* |
| 242 | *GGTCGCGGCGATAAAGACA* |
| 243_ 2 | *GACGTAGTTCGTTCAGGCGAGCAG* |

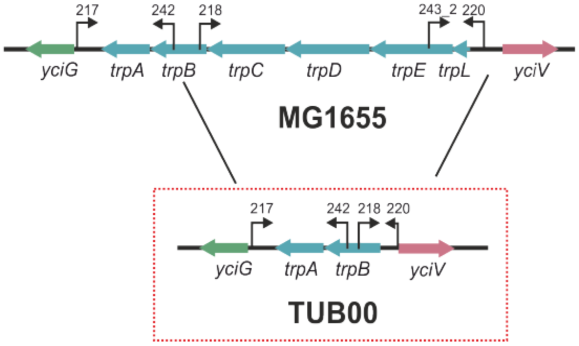

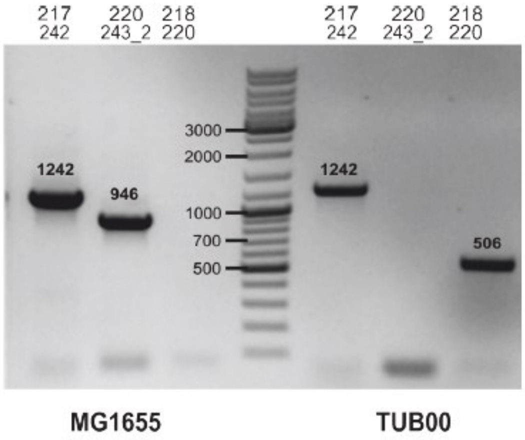

B

**Verification of phage assisted knockout of lysA**

| | |
|---|---|
| C1 | *CTCTCGCAATCCGGTAATCC* |
| C2 | *GCTGGTACGTCGTCATTCAG* |
| C5 | *AATGTGCATGTGAATGCCG* |

**Verification of phage assisted knockout of argA**

| | |
|---|---|
| C1 | *TAAACGACGACGGCTGATTG* |
| C2 | *CTGGCGGCGTTGTTTAGTTC* |
| C5 | *CATTAGTGACGCCCTGGGAA* |

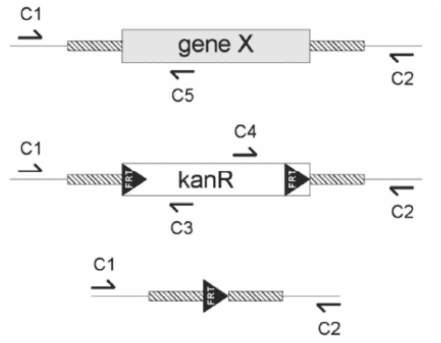

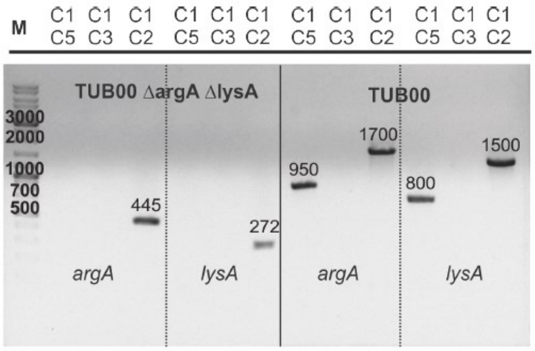

**Figure S1 I Strain construction.** A) Knockout of Trp biosynthesis pathway for ALE. Top left: Primers used for the verification. Top right: Schematic overview of MG1655 and TUB00 strain compositions. Bottom: Agarose gel demonstrating the successful KO by comparing MG1655 (wt) to TUB00. B) Knockout of *lysA* and *argA* for SILAC. Top left: Primers used for the verification. Top right: Schematic overview of where the primers bind. Bottom: Agarose gel showing successful KO by comparing TUB00 to the SILAC KO strain.

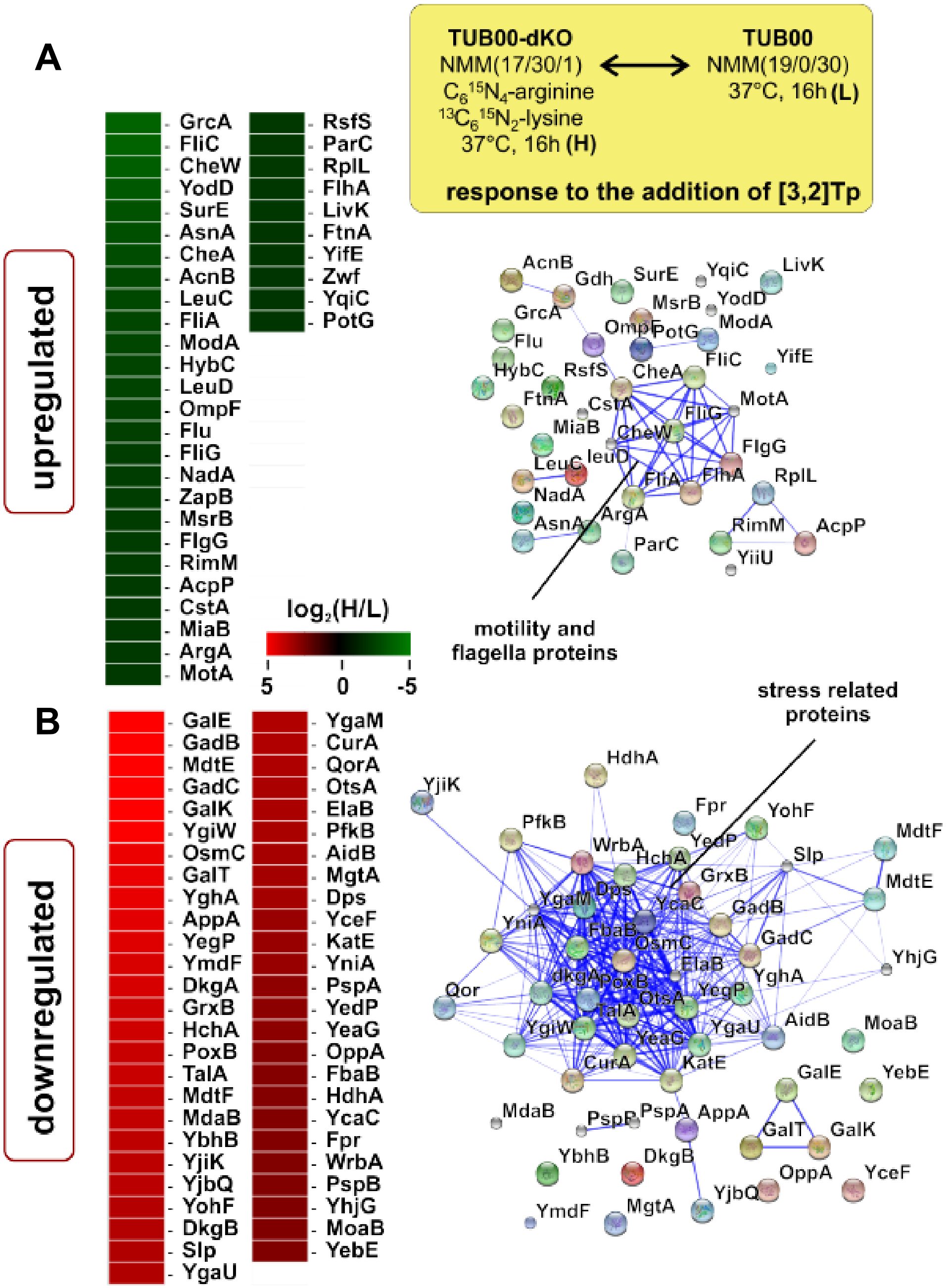

**Figure S2 I SILAC: Response of TUB00 to the addition of [3,2]Tp.** The heat map representation illustrates the 87 most A) up- and B) downregulated proteins as a response of TUB00 to the addition of [3,2]Tp (30 µM) in a limiting indole (1 µM) environment. The normalized H/L ratios (log2-transformed) are plotted of 'light'-labeled (L) TUB00 (NMM(19/0/30) and 'heavy'-labeled (H) TUB00dKO (NMM(17/30/1) + $^{13}C_6{}^{15}N_4$-arginine and $^{13}C_6{}^{15}N_2$-lysine). The cluster plot was created with string.db and illustrates protein-protein interaction networks.

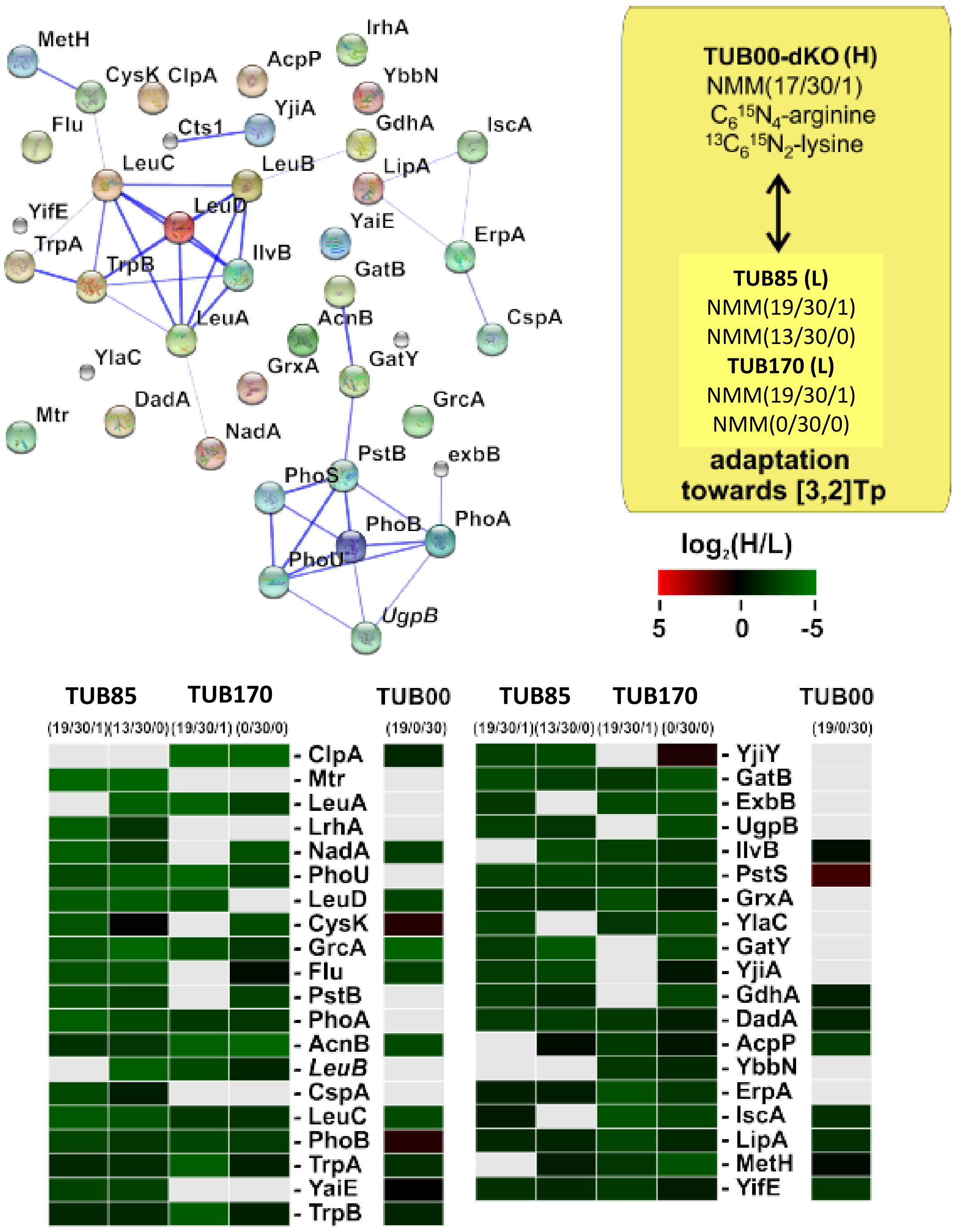

**Figure S3 I SILAC: The most upregulated proteins of the evolvates TUB85 and TUB170 in response to [3,2]Tp-supplementation.** The quantitative SILAC-based protein expression results shown above indicate the change in the regulatory network of the evolved derivatives TUB85 and TUB170 in comparison to the ancestral strain TUB00. Shown here are the most upregulated proteins. To rule out medium effects, the strains were grown in both indole-containing NMM(19/30/1) media and in their respective stringent indole-deficient NMM(19/30/0) media. In the column to the right of the protein names, the respective results for TUB00 grown in NMM(19/0/30) are shown. The normalized H/L ratios (log2-transformed) are plotted, in all cases 'heavy'-labeled (H) TUB00dKO cultivated in NMM(17/30/1) + $^{13}C_6{}^{15}N_4$-arginine and $^{13}C_6{}^{15}N_2$-lysine was used as the standard, whereas TUB00, TUB85 and TUB170 were 'light'-labeled (L) in their respective media. The cluster plot in the top left was created with string.db and illustrates protein-protein interaction networks.

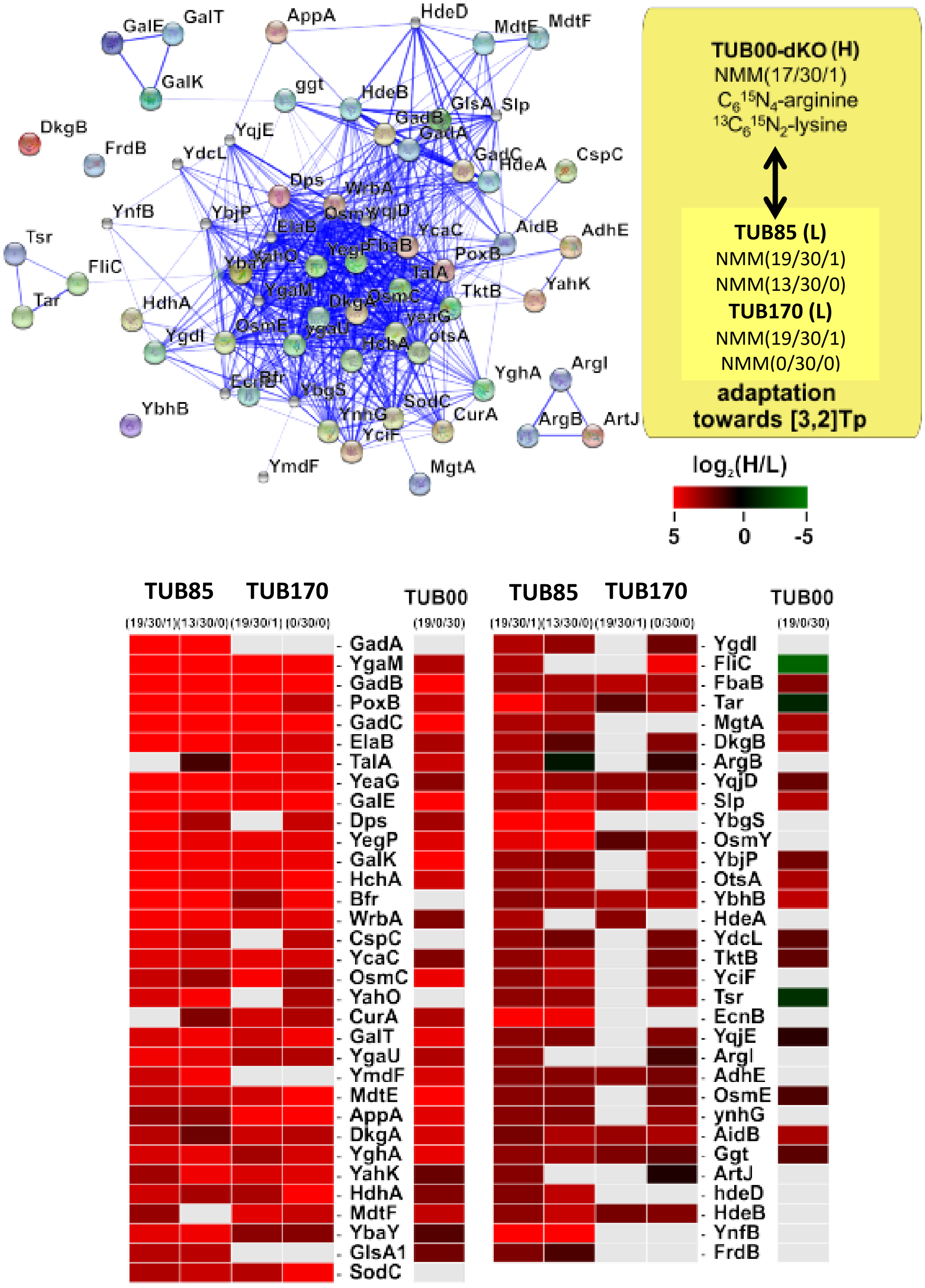

**Figure S4 I SILAC: The most downregulated proteins of the evolvates TUB85 and TUB170 in response to [3,2]Tp-supplementation.** The quantitative SILAC-based protein expression results shown above indicate the change in the regulatory network of the evolved derivatives TUB85 and TUB170 in comparison to the ancestral strain TUB00. To rule out medium effects, the strains were grown in both indole-containing NMM(19/30/1) media and in their respective stringent indole-deficient NMM(19/30/0) media. In the column to the right of the protein names, the respective results for TUB00 grown in NMM(19/0/30) are shown. The normalized H/L ratios (log2-transformed) are plotted, in all cases 'heavy'-labeled (H) TUB00dKO cultivated in NMM(17/30/1) + $^{13}C_6{}^{15}N_4$-arginine and $^{13}C_6{}^{15}N_2$-lysine was used as the standard, whereas TUB00, TUB85 and TUB170 were 'light'-labeled (L) in their respective media. The cluster plot in the top left was created with string.db and illustrates protein-protein interaction networks.
